# Effects of taurine in mice and zebrafish behavioral assays with translational relevance to schizophrenia

**DOI:** 10.1101/2022.03.29.486302

**Authors:** Franciele Kich Giongo, Matheus Gallas-Lopes, Radharani Benvenutti, Adrieli Sachett, Leonardo Marensi Bastos, Adriane Ribeiro Rosa, Ana Paula Herrmann

**Affiliations:** Laboratório de Neurobiologia e Psicofarmacologia Experimental (PsychoLab), Departamento de Farmacologia, Instituto de Ciências Básicas da Saúde, Universidade Federal do Rio Grande do Sul, Av. Sarmento Leite 500, Porto Alegre, Rio Grande do Sul, 90050-170, Brazil; Programa de Pós-Graduação em Farmacologia e Terapêutica, Instituto de Ciências Básicas da Saúde, Universidade Federal do Rio Grande do Sul, Av. Sarmento Leite 500, Porto Alegre, Rio Grande do Sul, 90050-170, Brazil; Programa de Pós-Graduação em Neurociências, Instituto de Ciências Básicas da Saúde, Universidade Federal do Rio Grande do Sul, Av. Sarmento Leite 500, Porto Alegre, Rio Grande do Sul, 90050-170, Brazil; Laboratório de Psiquiatria Molecular, Hospital de Clínicas de Porto Alegre, Rua Ramiro Barcelos, 2400, Porto Alegre, Rio Grande do Sul, 90035-003, Brazil

**Keywords:** schizophrenia, taurine, MK-801, C57BL/6, zebrafish, behavior

## Abstract

**Background:** Altered redox state and developmental abnormalities in glutamatergic and GABAergic transmission during development are linked to the behavioral changes associated with schizophrenia. As an amino acid that exerts antioxidant and inhibitory actions in the brain, taurine is a potential candidate to modulate biological targets relevant to this disorder. Here, we investigated in mice and zebrafish assays whether taurine prevents the behavioral changes induced by acute administration of MK-801 (dizocilpine), a glutamate NMDA receptor antagonist.

**Methods:** C57BL/6 mice were intraperitoneally administered with saline or taurine (50, 100 and 200 mg/kg) followed by MK-801 (0.15 mg/kg). Locomotor activity, social interaction and prepulse inhibition of the acoustic startle reflex were then assessed in different sets of animals. Zebrafish were exposed to tank water or taurine (42, 150 and 400 mg/L) followed by MK-801 (5 μM); social interaction and locomotor activity were evaluated in the same test.

**Results:** MK-801 induced hyperlocomotion and disrupted sensorimotor gating in mice; in zebrafish, it reduced sociability while increased locomotion. Taurine was mostly devoid of effects and did not counteract NMDA antagonism in mice or zebrafish.

**Discussion:** Contradicting previous clinical and preclinical data, taurine did not show antipsychotic-like effects in the present study. However, it still warrants consideration as a preventive intervention in animal models of relevance to the prodromal phase of schizophrenia; further studies are thus necessary to evaluate whether and how taurine might benefit patients.

## 1. Introduction

Schizophrenia is a serious mental illness that remains as one of the main challenges in modern psychiatry. With an incidence slightly higher in men than women (Jongsma et al., 2019), schizophrenia affects approximately one in a hundred people (McGrath et al., 2008) and dramatically changes the individual’s life course. Psychotic symptoms, social isolation, cognitive impairment, and stigma are only a few of the obstacles that contribute to the poor prognosis of this condition. Hyperdopaminergic activity in subcortical areas is associated with the onset of psychotic symptoms (McCutcheon et al., 2018), and might be linked to the loss of fast-spiking parvalbumin-positive GABAergic interneurons, hypofunction of glutamate NMDA receptors and oxidative stress (Cabungcal et al., 2013; Grace, 2016; Hardingham and Do, 2016). Furthermore, several studies have indicated increased frequency of smoking (Lasser et al., 2000), alcohol or illegal substances misuse (Hjorthøj et al., 2015), sedentary lifestyle (Stubbs et al., 2016) and poor dietary habits (Dipasquale et al., 2013) among individuals with schizophrenia, which results in a 13-15 years reduction in life expectancy, currently averaged at 65 years (Hjorthøj et al., 2017).

Early and continuous administration of antipsychotic drugs is the main pharmacological strategy; unfortunately, however, the currently available drugs do not provide a full recovery and a third of the patients are unresponsive to treatment (Jääskeläinen et al., 2013; Bruijnzeel et al., 2014). Antipsychotic drugs, which act by blocking dopamine D_2_ receptors, ameliorate mainly the positive symptoms (e.g., hallucinations and delusions), with little to no effect on negative symptoms (e.g., social isolation, depression, avolition) and cognitive impairment (e.g., poor long-term memory, sustained attention and cognitive performance) (Shafer and Dazzi, 2019). Pharmacological interventions pose yet another challenge: side effects such as extrapyramidal symptoms, metabolic syndrome and weight gain further impact quality of life and compromise adherence to treatment (Ellenbroek, 2012; Bruijnzeel et al., 2014).

Since the observation that phencyclidine induces in healthy individuals a psychotic state that resembles schizophrenia (Luisada and Brown, 1976; Allen and Young, 1978), NMDA antagonists have been used to recapitulate relevant behavioral alterations in animal models (Jones et al., 2011). Other non-competitive NMDA antagonists, such as MK-801 (dizocilpine) and ketamine, also trigger complex effects similar to the positive, negative and cognitive symptoms experienced by individuals with schizophrenia (Buffalo et al., 1994; Adler et al., 1999; Bondi et al., 2012). Here, we used acute administration of MK-801 in mice and zebrafish to study the antipsychotic-like properties of taurine in behavioral assays with translational relevance to schizophrenia.

Taurine, also known as 2-aminoethanesulfonic acid, is the most abundant free amino acid in the human body (Huxtable, 1992) and acts as an inhibitory neuromodulator in the brain (Oja and Saransaari, 1996); it also has neuroprotective, antioxidant and immunomodulatory properties (Almarghini et al., 1991; Redmond et al., 1998). The effects of taurine in the central nervous system are likely mediated by agonism at GABAergic and glycinergic receptors, yet it is still unknown whether taurine-specific receptors could be involved (Oja and Saransaari, 2015). Several studies have shown altered levels of taurine in the brain and plasma of schizophrenia patients and in animal models. Increased taurine levels were observed in the prefrontal cortex of schizophrenia patients, as well as a correlation between increased taurine and disease duration (Shirayama et al., 2010). A recent study also observed elevated taurine levels in serum samples of first psychotic episode and early stage patients (Parksepp et al., 2020). In contrast, decreased taurine levels were observed in the cerebrospinal fluid of drug-naïve individuals (Do et al., 1995). In a mice model of prenatal immune activation, taurine was decreased in the hippocampus, striatum, temporal and parietal cortex (Winter et al., 2009; Yang et al., 2019). Finally, a recent double-blind randomized trial on 86 individuals with first-episode psychosis showed a significant improvement in schizophrenia symptoms in patients who received taurine as an adjuvant treatment as compared to placebo (O’Donnell et al., 2016), while preclinical zebrafish data further support the potential benefits of taurine in this context (Franscescon et al., 2020, 2021).

Considering the above-cited evidence, this study aimed to test the hypothesis that taurine prevents schizophrenia-relevant behavioral alterations induced by acute administration of MK-801 in mice and zebrafish assays.

## 2. Materials and methods

### 2.1. Animals

#### 2.1.1. Mice

C57BL/6 male mice (7 to 14-week-old, 20-30 g) were obtained from an external vendor (Centro de Cardiologia Experimental – Instituto de Cardiologia, RS, Brazil). Upon arrival at Unidade de Experimentação Animal (Hospital de Clínicas de Porto Alegre), animals were housed in groups of 3-5 animals per cage (20 × 30 × 13 cm) for at least two weeks before experiments. Different sets of animals were used for each of the behavioral assays. The animals were maintained under controlled environmental conditions (reversed 12-h light/dark cycle with lights on at 7:00 a.m. and constant temperature of 22 ± 1 °C) with free access to food (Nuvilab CR-1^®^, PR, Brazil) and water. All procedures were approved by the animal welfare and ethical review committee of Hospital de Clínicas de Porto Alegre (approval #180498) and were performed in accordance with the relevant guidelines on care and use of laboratory animals and the Brazilian legislation.

#### 2.1.2. Zebrafish

Experiments were performed using male and female (50:50 ratio) short-fin wild-type zebrafish (6-month-old, 400-500 mg). Adult animals were obtained from a local commercial supplier (Delphis, RS, Brazil). The animals were maintained at Instituto de Ciências Básicas da Saúde in a light/dark cycle of 14/10 h with lights on at 7:00 a.m for at least two weeks before tests. Fish were kept in 16-L (40 × 20 × 24 cm) unenriched glass tanks with nonchlorinated water at a maximum density of two animals per liter. Tank water satisfied the controlled conditions required for zebrafish (26 ± 2 °C; pH 7.0 ± 0.3; dissolved oxygen at 7.0 ± 0.4 mg/L; total ammonia at <0.01 mg/L; total hardness at 5.8 mg/L; alkalinity at 22 mg/L CaCO_3_; conductivity of 1,500–1,600 μS/cm) and was constantly filtered by mechanical, biological, and chemical filtration systems (Altamar^®^, SP, Brazil). Food was provided twice a day as commercial flake food (Poytara^®^, SP, Brazil) plus brine shrimp (*Artemia salina*). The sex of the animals was confirmed after euthanasia by dissecting and analyzing the gonads. Animals were euthanized by hypothermic shock according to the AVMA Guidelines for the Euthanasia of Animals (Leary and Johnson, 2020). For all experiments, no sex effects were observed, so data were pooled together. All procedures were approved by the animal welfare and ethical review committee at the Universidade Federal do Rio Grande do Sul (approval #35525).

### 2.2. Drugs

Taurine and MK-801 (dizocilpine) were purchased from Sigma-Aldrich (St. Louis, MO, USA). For rodent experiments, drugs were dissolved in saline (0.9% NaCl) and solutions were freshly prepared and injected intraperitoneally (i.p.) at a volume of 5 mL/kg. Animals were manually contained for drug administration. For the zebrafish assay, MK-801 and taurine were dissolved in tank water; solutions were freshly prepared and renovated halfway through the experiment.

### 2.3. Experimental design

Different sets of animals were used for each experiment, totaling 288 mice and 96 zebrafish in the study. The animals were allocated to the experimental groups following block randomization procedures to counterbalance for litter and cage in mice experiments, and sex and home tank in zebrafish experiments. The order for outcome assessment was also randomized and care was taken to counterbalance the test apparatuses across the experimental groups. Outcome assessors were blind to the experimental groups, as well as the experimenters responsible for taking the animal and placing it in the test apparatus. An overview of the experimental design is illustrated in Figure 1.

**Figure 1.**
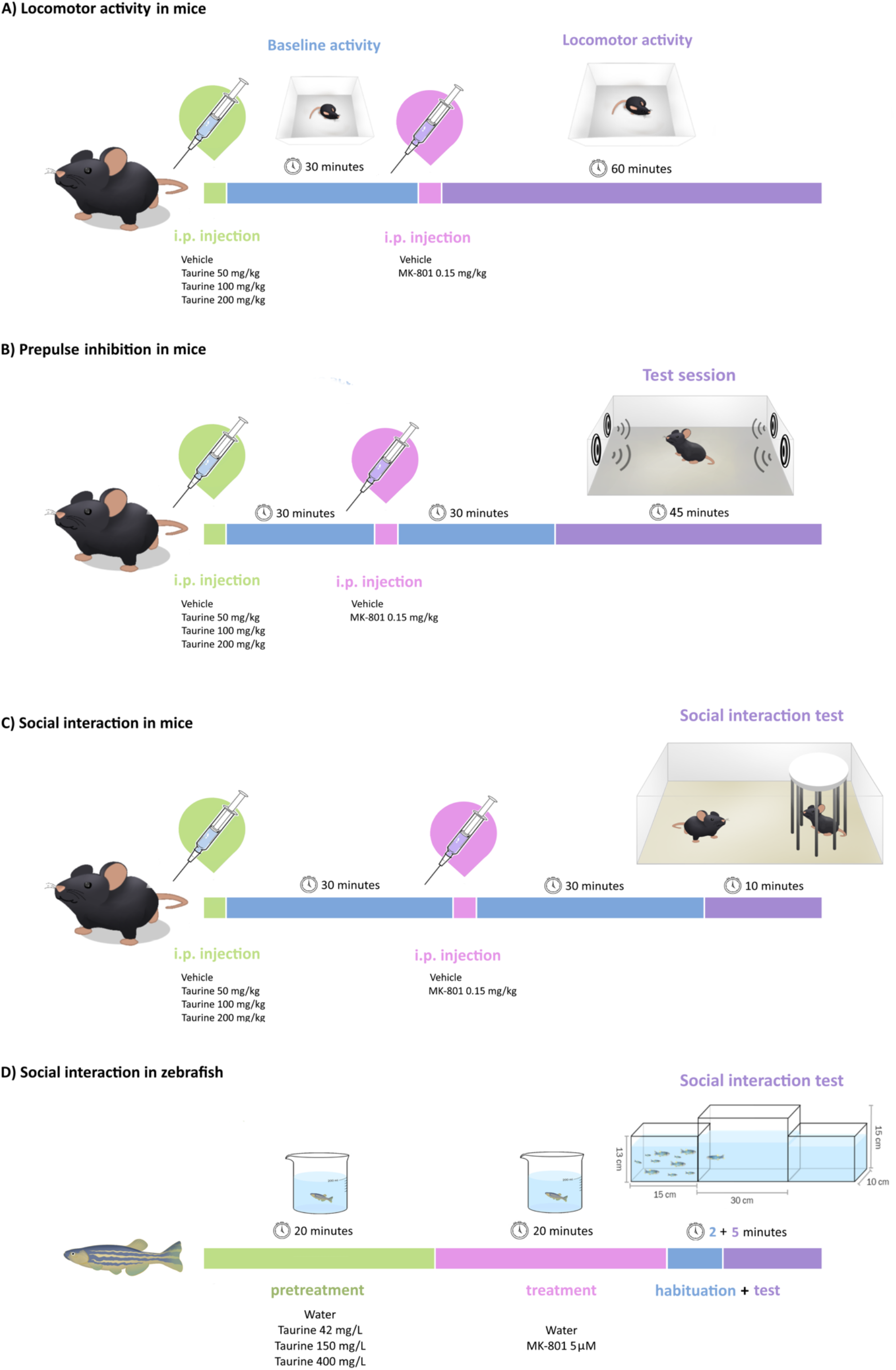
Overview of the experimental design. (A) Locomotor activity in mice, (B) prepulse inhibition of the startle reflex in mice, (C) social interaction in mice, (D) social interaction in zebrafish.

We report how we determined our sample size, all data exclusions, all manipulations, and all measures in the study. Raw data and analyses outputs were deposited in the Open Science Framework and are openly available at https://osf.io/qy2uw (Giongo et al., 2022).

#### 2.3.1. Locomotor activity in the open field in mice

The protocol for assessing the locomotor response to MK-801 was adapted from Meyer et al. (2008). Mice were injected with saline (0.9% NaCl) or taurine (50, 100, or 200 mg/kg, i.p.) and then placed in the center of an open field arena (40 × 40 × 40 cm) for baseline activity measurement during 30 min. Animals were then briefly removed to receive an i.p. injection of MK-801 (0.15 mg/kg); they were returned to the open field and locomotor activity was recorded for 60 min. The distance traveled in segments of 5 min was automatically scored using ANY-Maze software (Stoelting Co., Wood Dale, IL, USA).

#### 2.3.2. Prepulse inhibition of the acoustic startle reflex in mice

Sensorimotor gating was assessed by measuring the prepulse inhibition (PPI) of the acoustic startle reflex, which refers to the attenuation of the reaction to a startling stimulus (pulse) when it is shortly preceded by a weaker stimulus (prepulse). The protocol was adapted based on the methodology fully described elsewhere (Meyer et al., 2005). The apparatus consisted of two sound-attenuated startle chambers (Insight^®^, SP, Brazil) equipped with a movement-sensitive platform above which an acrylic enclosure was placed. One day prior to the experiment, subjects were habituated for 10 min to the apparatus with background noise (65 dB_A_). On the experiment day, animals received an i.p. injection of either saline (0.9% NaCl) or taurine (50, 100, or 200 mg/kg), followed 30 min later by an i.p. injection of saline or MK-801 (0.15 mg/kg). Testing started 30 min after the last drug administration. During a 45-min session, animals were presented to a series of stimuli comprising a mixture of four trial types: pulse-alone, prepulse-plus-pulse, prepulse-alone and no-stimulus (background noise, 65 dB_A_). The startle program consisted of three different intensities of a 40-ms white noise pulse (100, 110, and 120 dB_A_) combined or not with three different intensities of a 20-ms prepulse (71, 77, and 83 dB_A_, which corresponded to 6, 12, and 18 dB above background, respectively). The stimulus-onset asynchrony of the prepulse and pulse stimuli on all prepulse-plus-pulse trials was 100 ms (onset-to-onset). Each session began with a 2-min acclimation period in the enclosure, followed by 6 consecutive pulse-alone trials to habituate and stabilize the startle response. Subsequently, each stimulus was presented 12 times in a pseudorandom order with an average interval between successive trials of 15 ± 5 s. The session was concluded with 6 consecutive pulse-alone trials. Boxes were cleaned with water and dried between sessions. For each subject, PPI was indexed as mean percent inhibition of startle response obtained in the prepulseplus-trials compared to pulse-alone trials by following the expression: [1-(mean reactivity on prepulse-plus-pulse trials / mean reactivity on pulse-alone trials) × 1/100]. The first and last six trials were not included in the calculation of percent PPI. In addition to PPI, reactivity to prepulse- and pulse-alone trials were also analyzed.

#### 2.3.3. Social interaction in mice

The social interaction protocol was adapted from Jeevakumar et al. (2015). Experimental and stimulus mice were isolated for 24 h prior to testing. On the day of the experiment, mice received an i.p. injection of either saline (0.9% NaCl) or taurine (50, 100, or 200 mg/kg) followed by another i.p. injection of saline or MK-801 (0.15 mg/kg) 30 min later. Testing began 30 min after the last injection. A stimulus mouse was placed inside a cylindrical custom-built container (20 cm high, steel bars separated by 1 cm, acrylic lid) and then introduced to the home cage of experimental mice for 10 min. All sessions were video-recorded and interaction time (defined as sniffing and investigating at close proximity) were scored offline using Behavioral Observation Research Interactive Software (BORIS; Friard & Gamba, 2016).

#### 2.3.4. Social interaction in zebrafish

The protocol for the social interaction test in zebrafish followed the method described by Benvenutti et al. (2021). Animals were individually exposed to water or taurine solutions at 42, 150 or 400 mg/L in 500-mL beakers containing 200-mL solution for 20 min. They were then transferred to another beaker containing either water or MK-801 at 5 μM for another 20 minutes. After exposure, animals were placed for 7 min in a tank (30 × 10 × 15 cm) flanked by two identical tanks (15 × 10 × 13 cm) either empty (neutral stimulus) or containing 10 unknown zebrafish (social stimulus). All three tanks were filled with water in standard conditions at a level of 10 cm. The position of the social stimulus (right or left) was counterbalanced throughout the tests. The water in the test tanks was changed between every animal. To assess social behavior, the test apparatus was virtually divided into three vertical zones (interaction, middle, and neutral). Animals were habituated to the apparatus for 2 min and then analyzed for the last 5 min. Videos were recorded from the front view and time spent in the interaction zone was quantified as a proxy for social interaction. Additionally, total distance traveled, number of crossings between the vertical zones of the tank and immobility time were quantified as secondary locomotor parameters. All outcomes were automatically scored using ANY-Maze software (Stoelting Co., Wood Dale, IL, USA).

### 2.4. Statistical analysis

Outliers were defined following the rule of mean ± 2 standard deviations. This resulted in five outliers removed from the PPI test (one TAU 50/CTRL, one TAU 100/CTRL, two TAU 50/MK and one TAU 100/MK), five outliers removed from the social interaction test in mice (one TAU 50/CTRL, one TAU 100/CTRL, one TAU 200/CTRL, one TAU 50/MK and one TAU 200/MK), and two outliers removed from the social interaction test in zebrafish (one CTRL/CTRL and one TAU 42/CTRL). No outliers were removed from the hyperlocomotion test (mean distance traveled after MK-801 was the outcome used for the check). Two mice from the social interaction set died of unknown causes after allocation but before testing (one CTRL/MK and one TAU 100/MK).

The sample size to detect a 0.5 effect size with 0.95 power and 0.05 alpha was calculated using Minitab (version 21.1) for Windows; this resulted in n=10 for locomotor activity (4 groups) and n=12 for all other assays (8 groups). GraphPad Prism 8 (version 8.4.3) for macOS was used to run the statistical analyses and plot the results. For locomotor activity, distance traveled as a function of 5-min time segments was analyzed using repeated measures ANOVA, with time as the within-subjects factor, and taurine pretreatment as the between-subjects factor; the two phases of the experiment (i.e., baseline and MK-801 treatment) were analyzed separately. The data from the remaining experiments were analyzed by two-way ANOVA, with taurine pretreatment and MK-801 treatment as the main factors. Bonferroni post hoc test was applied as appropriate. The significance level was set at p<0.05. Data were expressed as mean ± standard deviation.

## 3. RESULTS

### 3.1. Locomotor activity in response to MK-801

The hyperlocomotion in response to an MK-801 challenge was assessed as an outcome related to the positive symptoms of schizophrenia (Powell and Geyer, 2007). Figure 2A shows that distance travelled by the mice in the open field increased after the MK-801 challenge in all experimental groups (time effect: F_11,396_ = 136.3, p<0.0001). Taurine did not prevent the effects of MK-801 (taurine effect: F_3,36_ = 1.471, p=0.2387; interaction effect: F_33,396_ = 0.9391, p=0.5675). In the baseline phase (first 30 min), locomotion decreased as animals habituated to the environment (time effect: F_5,180_ = 102.0, p<0.0001), and no differences between the groups were observed (taurine effect: F_3,36_ = 0.1631, p=0.9205; interaction effect: F_15,180_ = 1.140, p=0.3239).

**Figure 2.**
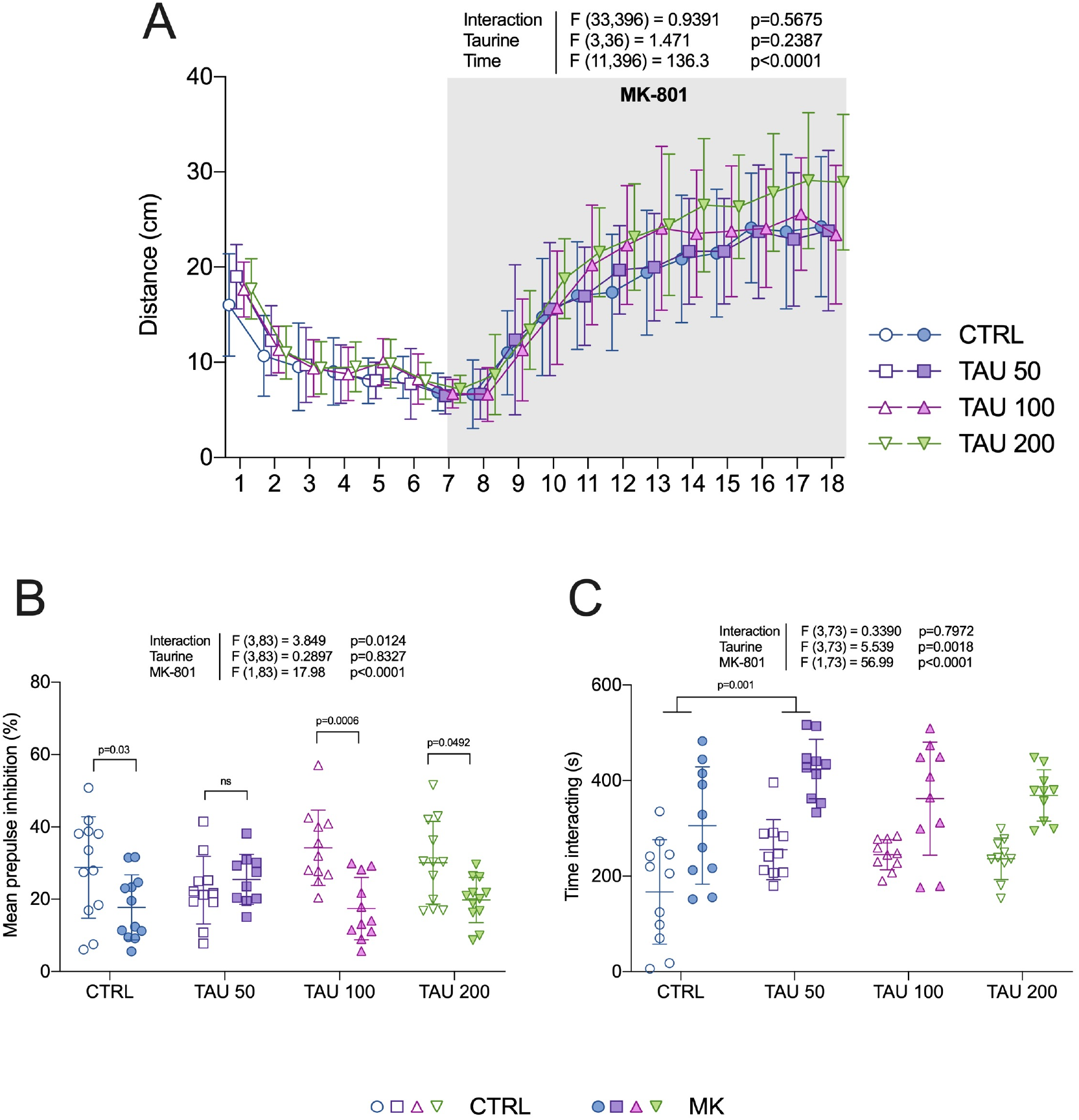
Effects of taurine on behavioral abnormalities induced by MK-801 in mice. (A) Locomotor activity (in segments of 5 min), (B) prepulse inhibition of the startle reflex and (C) social interaction were evaluated as measures relevant to the positive, cognitive, and negative symptoms of schizophrenia, respectively. Two-way ANOVA followed by Bonferroni post hoc test. Data are presented as mean ± standard deviation. n=10-12. CTRL: control, TAU: taurine (doses are denoted in mg/kg).

### 3.2. MK-801-induced prepulse inhibition deficits

The effects of MK-801 on sensorimotor gating were assessed by the paradigm of prepulse inhibition (PPI) of the acoustic startle reflex, which is a translational measure related to the cognitive symptoms of schizophrenia (Powell et al., 2009). Mice treated with MK-801 showed lower levels of PPI when compared to controls (MK-801 main effect: F_1,83_ = 17.98, p<0.0001), indicating deficits in sensorimotor gating (Figure 2B). Two-way ANOVA also revealed a significant interaction effect (F_3,83_ = 3.849, p=0.0124) in the absence of a taurine main effect (F_3,83_ = 0.2897, p=0.8327). Bonferroni post hoc tests comparing MK-801 groups to their respective controls resulted in significant differences for all comparisons except for the groups pretreated with taurine at 50 mg/kg (lowest dose), indicating an attenuation of the PPI deficit induced by MK-801.

### 3.3. Social interaction in mice

The social behavior towards an unfamiliar mouse introduced in the home cage was evaluated as a phenotype relevant to the negative symptoms of schizophrenia (Jones et al., 2011). Subject mice were manually scored according to their interest in sniffing or investigating at proximity the enclosed stimulus mouse. Figure 2C shows that groups treated with MK-801 spent more time interacting with the social stimulus (MK-801 main effect: F_1,73_ = 56.99, p<0.0001). Two-way ANOVA also revealed a taurine main effect (F_3,73_ = 5.539, p=0.0018) without a significant interaction (F_3,73_ = 0.339, p=0.7972). Bonferroni post hoc comparisons restricted to the pretreatment factor (taurine main effect) indicated that interaction time was significantly higher in groups pretreated with taurine at 50 mg/kg in comparison to control groups (p=0.001).

Data are presented as mean ± SD. n=11-12. CTRL: control, TAU: taurine (exposure concentrations are denoted in mg/L).

### 3.4. Social interaction in zebrafish

Zebrafish is increasingly considered as a model organism suitable to study drug-induced behavioral phenotypes relevant to schizophrenia (Gawel et al., 2019; Benvenutti et al., 2021). Figure 3A shows that zebrafish exposed to MK-801 spent less time in the side of the tank where conspecifics were presented, denoting decreased social preference in comparison to control groups (MK-801 main effect: F_1,86_ = 29.28, p<0.0001). The other outcomes evaluated in this test were also altered by MK-801, as shown by increases in total distance travelled (F_1,86_ = 5.74, p=0.0188; Fig. 3B), number crossings between the zones of the tank (F_1,86_ = 37.13, p<0.0001; Fig. 3C) and immobility time (F_1,86_ = 8.597, p=0.0043; Fig. 3D). Taurine was devoid of effects in all parameters, as no main effects for drug pretreatment or interaction effects were observed.

**Figure 3.**
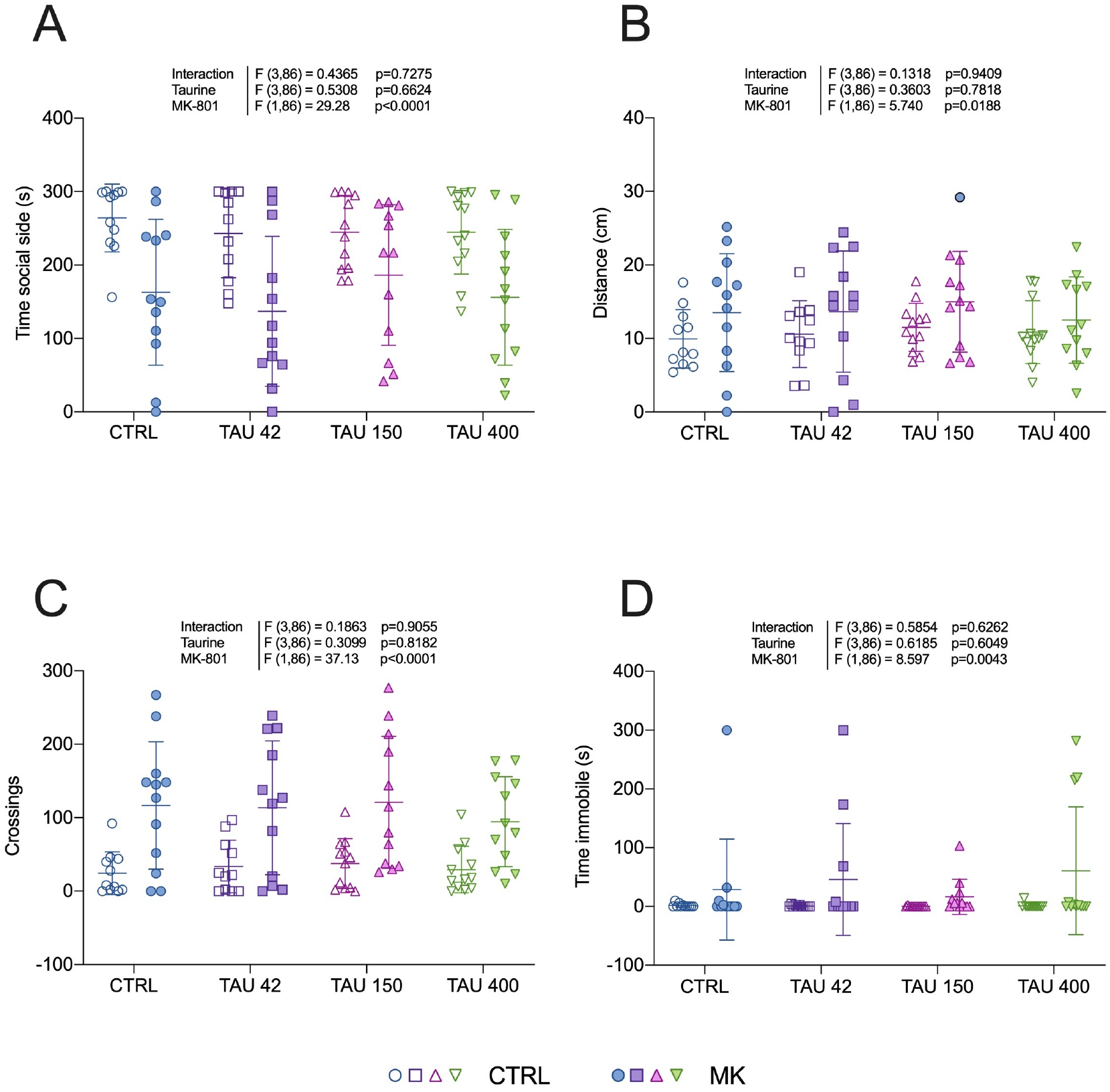
Effects of taurine in the social interaction test in zebrafish. (A) Time spent in the social stimulus side was measured as a proxy for social interaction, while (B) total distance traveled, (C) number of crossings and (D) time spent immobile were quantified as secondary locomotor parameters. Two-way ANOVA.

## 4. DISCUSSION

Animal models provide a unique opportunity to understand how genetic, molecular, and environmental factors might lead to the development of schizophrenia. In this study, we used MK-801 to acutely induce behavioral alterations of translational relevance to schizophrenia in C57BL/6 mice and zebrafish, and ultimately evaluate the preventive effects of taurine against the deficits caused by NMDA antagonism in a two-species approach.

Taurine’s role has been studied in several conditions, including depression, cardiac failure, retina degeneration and growth problems (Lourenço and Camilo, 2002). Here, we found that taurine largely failed to prevent MK-801-induced behavioral alterations. Although a significant interaction was found in the prepulse inhibition (PPI) test and post hoc analysis showed that the group pretreated with taurine at 50 mg/kg before MK-801 administration was not significantly different than its respective control, this should be interpreted with caution as taurine at this dose seems to buffer PPI to intermediate levels instead of fully preventing the deficit induced by MK-801. In the social interaction test in mice, both groups treated with taurine at 50 mg/kg spent more time interacting when compared to pretreatment controls. This agrees with a previous study in which taurine at a similar dose (42 mg/kg) was shown to increase social interaction in Wistar rats (Kong et al., 2006); such increases in social interaction may be explained by the anxiolytic effects reported for taurine in several studies (Kong et al., 2006; El Idrissi et al., 2009; Mezzomo et al., 2016, 2019; Jung and Kim, 2019; Neuwirth et al., 2019; Fontana et al., 2020).

As expected, MK-801 increased the total distance traveled and caused a PPI deficit in mice. In zebrafish, we observed reduced time in the social side as well as hyperlocomotion in all groups exposed to MK-801. Curiously, mice treated with MK-801 spent more time interacting with the stimulus mice when compared to controls. This was unexpected since in most studies NMDA antagonism leads to decreased levels of social interaction (Morales and Spear, 2014; Zoicas and Kornhuber, 2019). Jeevakumar *et al*. (2015), for example, used a similar home cage protocol and observed a significantly reduced investigation time in adult mice exposed to ketamine in the second postnatal week. Although both ketamine and MK-801 are NMDA antagonists, differences might be related to MK-801 being a more specific NMDA antagonist, while ketamine also interacts with dopaminergic and serotoninergic systems (Kapur and Seeman, 2002; Stone et al., 2007). Drug administration regimen and protocol adaptations also might contribute to this difference in social behavior. Moreover, MK-801 showed a fast-acting but nonsustainable antidepressant response in control mice (Autry et al., 2011; Zanos et al., 2016), which could explain the increased social behavior in our experiment as mice were tested 30 minutes after the MK-801 injection.

It is well known that excessive stimulation of glutamatergic receptors causes excitotoxicity due to increased intracellular levels of calcium. Previous studies have demonstrated that taurine may act as a neuroprotector either by decreasing intracellular free calcium or by counterbalancing glutamatergic transmission via voltage-gated calcium channels (Lidsky et al., 1995; El Idrissi and Trenkner, 1999; Saransaari and Oja, 2000). Acamprosate, a synthetic analog of taurine, is hypothesized to decrease NMDA receptor activity by modulating the expression of NMDA receptor subunits in specific brain regions (Rammes et al., 2001; Heilig and Egli, 2006). In addition, Chan et al (2015) showed that taurine binds to GluN2B subunit of the NMDA receptor and causes a prolonged inhibition of excitatory synaptic transmission in an *ex vivo* model.

Although taurine was not able to prevent the deficits observed in our study, it still might have beneficial effects in psychosis models that better mimic the course of schizophrenia, such as neurodevelopmental models. Various studies linked behavioral and neurobiological dysfunctions of schizophrenia to neurodevelopment, which translates into symptoms that appear mainly during late adolescence (Brown, 2006; Knuesel et al., 2014; Volk and Lewis, 2014; Hantsoo et al., 2019). Interventions that aim to act in the prodrome period, preventing the first psychotic episode, are thought to have better outcomes than antipsychotic treatment, once they have been ultimately unsuccessful in preventing disease onset in individuals with schizophrenia (McGlashan et al., 2003; McGorry et al., 2013; Woods et al., 2017). Therefore, continuous taurine administration in vulnerability periods might prevent the abnormalities that emerge in early adulthood. With antioxidant and neuroprotector properties, taurine might be able to normalize the altered redox state and parvalbumin-positive interneurons loss found in animal models and patients with schizophrenia (Fung et al., 2010; Gill and Grace, 2014; Salim, 2014; Steullet et al., 2017; Kaar et al., 2019; Goh et al., 2022). Grace *et al*. (2016) hypothesized that the dysfunction of the dopaminergic system might be a consequence of the loss of a large number of fast-spiking parvalbumin-positive GABAergic interneurons in the ventral subiculum of the hippocampus, causing hyperactivation and dysrhythmic behavior of pyramidal neurons. Taurine might ameliorate this hyperactivation by compensating the inhibitory loss at parvalbumin-positive interneurons and preventing the onset of symptoms in a neurodevelopmental model. Since this dysregulation is postulated to occur in late adolescence or early adulthood, taurine should be administered prior to this period to prevent the disruption of basolateral amygdala, nucleus accumbens and prefrontal cortex activity and rhythmicity, all of which participate in circuits interconnected with the ventral subiculum.

In regards to zebrafish, it has been demonstrated that MK-801 induces hyperlocomotion, although it is not clear which neuronal mechanisms might be involved (Menezes et al., 2015; Tran et al., 2016; Benvenutti et al., 2021; Franscescon et al., 2021). Zebrafish increasing use is a great solution to avoid species biases and focus on a robust cross-species approach, aside from being an accessible way to screen for potential novels treatments (Bruni et al., 2016; Burrows and Hannan, 2016; Gawel et al., 2019). Taurine has been demonstrated to prevent MK-801 hyperlocomotion and memory impairment in zebrafish (Franscescon et al., 2020, 2021), a finding that we could not replicate in our study. Here, taurine was not able to counteract MK-801 effects on locomotor activity or social interaction. In our protocol, zebrafish were exposed to taurine and MK-801 in a beaker for 20 minutes, and time spent on the social side and distance traveled were assessed. Differences in drug administration route and exposure time might contribute to the divergent outcomes. Considering that we also observed a lack of a clear antipsychotic effect of taurine in rodents, we reckon that our findings are robust and consistent across species, which does not necessarily rule out taurine antipsychotic effect in other treatment regimens.

A limitation of our study is that it remains to be established whether taurine can prevent the neuropathological events of schizophrenia in preclinical models that better simulate the course of the disease. Our study was not designed to act in the prodromal phase of schizophrenia, which we believe is a key opportunity window to prevent the alterations that emerge in early adulthood. Another limitation is that preclinical models of schizophrenia likely do not reflect the neurobiology underlying the positive symptoms of the disease, making it difficult to be accurately assessed in behavior tests (Kesby et al., 2018). Because hallucinations are false percepts perceived subjectively as true, a valid assessment of this behavior in rodents can require extensive training, and thus are incompatible with time-sensitive analysis, such as MK-801 acute administration (Schmack et al., 2021). Therefore, other rodent preclinical models of schizophrenia that have a more long-lasting endophenotype, and wherefore allow this type of assessment, may appraise positive-like symptoms with a better predictive validity than the locomotory response to MK-801.

The frequent failure in translating preclinical findings to clinical settings has been increasingly discussed, and strategies to overcome this loss in translation have been suggested (Seyhan, 2019). The strength of our study lies in including two model organisms from different phylogenetic classes, which increases the external validity of preclinical studies. Though more studies are necessary to evaluate taurine’s role in schizophrenia, our two-species approach contradicts previous studies by showing that, at least acutely, taurine is not able to prevent the behavioral alterations induced by antagonism of NMDA receptors.

## 5. ACKNOWLEDGMENTS

This work was supported by Fundação de Amparo à Pesquisa do Estado do Rio Grande do Sul (FAPERGS), grant agreement number 19/2551-0001216-0, Pró-Reitoria de Pesquisa – Universidade Federal do Rio Grande do Sul (PROPESQ-UFRGS), and Fundo de Incentivo à Pesquisa e Eventos – Hospital de Clínicas de Porto Alegre (FIPE-HCPA). Fellowships were granted to F.G. and R.B. from Coordenação de Aperfeiçoamento de Pessoal de Nível Superior (CAPES), and to A.S. from Conselho Nacional de Desenvolvimento Científico e Tecnológico (CNPq). We would like to give a special thanks to Marta Cioato and the staff at Unidade de Experimentação Animal - Hospital de Clínicas de Porto Alegre (HCPA) for all the help provided.

## REFERENCES

Adler CM, Malhotra AK, Elman I, Goldberg T, Egan M, Pickar D, Breier A (1999) Comparison of ketamine-induced thought disorder in healthy volunteers and thought disorder in schizophrenia. Am J Psychiatry 156:1646–1649.

Allen RM, Young SJ (1978) Phencyclidine-induced psychosis. Am J Psychiatry 135:1081–1084.

Almarghini K, Remy A, Tappaz M (1991) Immunocytochemistry of the taurine biosynthesis enzyme, cysteine sulfinate decarboxylase, in the cerebellum: evidence for a glial localization. Neuroscience 43:111–119.

Autry AE, Adachi M, Nosyreva E, Na ES, Los MF, Cheng P, Kavalali ET, Monteggia LM (2011) NMDA receptor blockade at rest triggers rapid behavioural antidepressant responses. Nature 475:91–95.

Benvenutti R, Gallas-Lopes M, Sachett A, Marcon M, Strogulski NR, Reis CG, Chitolina R, Piato A, Herrmann AP (2021) How do zebrafish (*Danio rerio*) respond to MK-801 and amphetamine? Relevance for assessing schizophrenia-related endophenotypes in alternative model organisms. J Neurosci Res 99:2844–2859.

Bondi C, Matthews M, Moghaddam B (2012) Glutamatergic animal models of schizophrenia. Curr Pharm Des 18:1593–1604.

Brown AS (2006) Prenatal infection as a risk factor for schizophrenia. Schizophr Bull 32:200–202.

Bruijnzeel D, Suryadevara U, Tandon R (2014) Antipsychotic treatment of schizophrenia: an update. Asian J Psychiatr 11:3–7.

Bruni G et al. (2016) Zebrafish behavioral profiling identifies multitarget antipsychotic-like compounds. Nat Chem Biol 12:559–566.

Buffalo EA, Gillam MP, Allen RR, Paule MG (1994) Acute behavioral effects of MK-801 in rhesus monkeys: assessment using an operant test battery. Pharmacol Biochem Behav 48:935–940.

Burrows EL, Hannan AJ (2016) Cognitive endophenotypes, gene-environment interactions and experience-dependent plasticity in animal models of schizophrenia. Biol Psychol 116:82–89.

Cabungcal J-H, Steullet P, Kraftsik R, Cuenod M, Do KQ (2013) Early-life insults impair parvalbumin interneurons via oxidative stress: reversal by N-acetylcysteine. Biol Psychiatry 73:574–582.

Chan CY, Singh I, Magnuson H, Zohaib M, Bakshi KP, Le François B, Anazco-Ayala A, Lee EJ, Tom A, YeeMon K, Ragnauth A, Friedman E, Banerjee SP (2015) Taurine Targets the GluN2b-Containing NMDA Receptor Subtype. Adv Exp Med Biol 803:531–544.

Dipasquale S, Pariante CM, Dazzan P, Aguglia E, McGuire P, Mondelli V (2013) The dietary pattern of patients with schizophrenia: a systematic review. J Psychiatr Res 47:197–207.

Do KQ, Lauer CJ, Schreiber W, Zollinger M, Gutteck-Amsler U, Cuénod M, Holsboer F (1995) gamma-Glutamylglutamine and taurine concentrations are decreased in the cerebrospinal fluid of drug-naive patients with schizophrenic disorders. J Neurochem 65:2652–2662.

El Idrissi A, Boukarrou L, Heany W, Malliaros G, Sangdee C, Neuwirth L (2009) Effects of taurine on anxiety-like and locomotor behavior of mice. Adv Exp Med Biol 643:207–215.

El Idrissi A, Trenkner E (1999) Growth factors and taurine protect against excitotoxicity by stabilizing calcium homeostasis and energy metabolism. J Neurosci 19:9459–9468.

Ellenbroek BA (2012) Psychopharmacological treatment of schizophrenia: what do we have, and what could we get? Neuropharmacology 62:1371–1380.

Fontana BD, Duarte T, Müller TE, Canzian J, Ziani PR, Mezzomo NJ, Parker MO, Rosemberg DB (2020) Concomitant taurine exposure counteracts ethanol-induced changes in locomotor and anxiety-like responses in zebrafish. Psychopharmacology (Berl) 237:735–743.

Franscescon F, Müller TE, Bertoncello KT, Rosemberg DB (2020) Neuroprotective role of taurine on MK-801-induced memory impairment and hyperlocomotion in zebrafish. Neurochem Int 135:104710.

Franscescon F, Souza TP, Müller TE, Michelotti P, Canzian J, Stefanello FV, Rosemberg DB (2021) Taurine prevents MK-801-induced shoal dispersion and altered cortisol responses in zebrafish. Prog Neuropsychopharmacol Biol Psychiatry 111:110399.

Friard O, Gamba M (2016) BORIS: A free, versatile open-source event-logging sofware for video/audio coding and live observations. Methods in Ecology and Evolution 7:1325–1330.

Fung SJ, Webster MJ, Sivagnanasundaram S, Duncan C, Elashoff M, Weickert CS (2010) Expression of interneuron markers in the dorsolateral prefrontal cortex of the developing human and in schizophrenia. Am J Psychiatry 167:1479–1488.

Gawel K, Banono NS, Michalak A, Esguerra CV (2019) A critical review of zebrafish schizophrenia models: Time for validation? Neurosci Biobehav Rev 107:6–22.

Gill KM, Grace AA (2014) Corresponding decrease in neuronal markers signals progressive parvalbumin neuron loss in MAM schizophrenia model. Int J Neuropsychopharmacol 17:1609–1619.

Giongo FK, Gallas-Lopes M, Benvenutti R, Sachett A, Bastos LM, Rosa AR, Herrmann AP (2022) Effects of taurine in preclinical behavioral assays relevant to schizophrenia. Available at: https://osf.io/qy2uw/ [Accessed March 29, 2022].

Goh XX, Tang PY, Tee SF (2022) Effects of antipsychotics on antioxidant defence system in patients with schizophrenia: A meta-analysis. Psychiatry Res 309:114429.

Grace AA (2016) Dysregulation of the dopamine system in the pathophysiology of schizophrenia and depression. Nat Rev Neurosci 17:524–532.

Hantsoo L, Kornfield S, Anguera MC, Epperson CN (2019) Inflammation: A Proposed Intermediary Between Maternal Stress and Offspring Neuropsychiatric Risk. Biol Psychiatry 85:97–106.

Hardingham GE, Do KQ (2016) Linking early-life NMDAR hypofunction and oxidative stress in schizophrenia pathogenesis. Nat Rev Neurosci 17:125–134.

Heilig M, Egli M (2006) Pharmacological treatment of alcohol dependence: target symptoms and target mechanisms. Pharmacol Ther 111:855–876.

Hjorthøj C, Østergaard MLD, Benros ME, Toftdahl NG, Erlangsen A, Andersen JT, Nordentoft M (2015) Association between alcohol and substance use disorders and all-cause and cause-specific mortality in schizophrenia, bipolar disorder, and unipolar depression: a nationwide, prospective, register-based study. Lancet Psychiatry 2:801–808.

Hjorthøj C, Stürup AE, McGrath JJ, Nordentoft M (2017) Years of potential life lost and life expectancy in schizophrenia: a systematic review and meta-analysis. The Lancet Psychiatry 4:295–301.

Huxtable RJ (1992) Physiological actions of taurine. Physiol Rev 72:101–163.

Jääskeläinen E, Juola P, Hirvonen N, McGrath JJ, Saha S, Isohanni M, Veijola J, Miettunen J (2013) A systematic review and meta-analysis of recovery in schizophrenia. Schizophr Bull 39:1296–1306.

Jeevakumar V, Driskill C, Paine A, Sobhanian M, Vakil H, Morris B, Ramos J, Kroener S (2015) Ketamine administration during the second postnatal week induces enduring schizophrenia-like behavioral symptoms and reduces parvalbumin expression in the medial prefrontal cortex of adult mice. Behav Brain Res 282:165–175.

Jones C, Watson D, Fone K (2011) Animal models of schizophrenia. Br J Pharmacol 164:1162–1194.

Jongsma HE, Turner C, Kirkbride JB, Jones PB (2019) International incidence of psychotic disorders, 2002-17: a systematic review and meta-analysis. Lancet Public Health 4:e229–e244.

Jung JH, Kim S-J (2019) Anxiolytic Action of Taurine via Intranasal Administration in Mice. Biomol Ther (Seoul) 27:450–456.

Kaar SJ, Angelescu I, Marques TR, Howes OD (2019) Pre-frontal parvalbumin interneurons in schizophrenia: a meta-analysis of post-mortem studies. J Neural Transm (Vienna) 126:1637–1651.

Kapur S, Seeman P (2002) NMDA receptor antagonists ketamine and PCP have direct effects on the dopamine D2 and serotonin 5-HT2 receptors—implications for models of schizophrenia. Mol Psychiatry 7:837–844.

Kesby JP, Eyles DW, McGrath JJ, Scott JG (2018) Dopamine, psychosis and schizophrenia: the widening gap between basic and clinical neuroscience. Transl Psychiatry 8:30.

Knuesel I, Chicha L, Britschgi M, Schobel SA, Bodmer M, Hellings JA, Toovey S, Prinssen EP (2014) Maternal immune activation and abnormal brain development across CNS disorders. Nat Rev Neurol 10:643–660.

Kong WX, Chen SW, Li YL, Zhang YJ, Wang R, Min L, Mi X (2006) Effects of taurine on rat behaviors in three anxiety models. Pharmacol Biochem Behav 83:271–276.

Lasser K, Boyd JW, Woolhandler S, Himmelstein DU, McCormick D, Bor DH (2000) Smoking and mental illness: A population-based prevalence study. JAMA 284:2606–2610.

Leary, Johnson (2020) AVMA guidelines for the euthanasia of animals: 2020 edition. Available at: https://www.avma.org/sites/default/files/2020-02/Guidelines-on-Euthanasia-2020.pdf [Accessed March 13, 2022].

Lidsky TI, Schneider JS, Yablonsky-Alter E, Zuck LG, Banerjee SP (1995) Taurine prevents haloperidol-induced changes in striatal neurochemistry and behavior. Brain Res 686:104–106.

Lourenço R, Camilo ME (2002) Taurine: a conditionally essential amino acid in humans? An overview in health and disease. Nutr Hosp 17:262–270.

Luisada PV, Brown BI (1976) Clinical management of the phencyclidine psychosis. Clin Toxicol 9:539–545.

McCutcheon R, Beck K, Jauhar S, Howes OD (2018) Defining the Locus of Dopaminergic Dysfunction in Schizophrenia: A Meta-analysis and Test of the Mesolimbic Hypothesis. Schizophr Bull 44:1301–1311.

McGlashan TH, Zipursky RB, Perkins D, Addington J, Miller TJ, Woods SW, Hawkins KA, Hoffman R, Lindborg S, Tohen M, Breier A (2003) The PRIME North America randomized double-blind clinical trial of olanzapine versus placebo in patients at risk of being prodromally symptomatic for psychosis. I. Study rationale and design. Schizophr Res 61:7–18.

McGorry PD, Nelson B, Phillips LJ, Yuen HP, Francey SM, Thampi A, Berger GE, Amminger GP, Simmons MB, Kelly D, Dip G, Thompson AD, Yung AR (2013) Randomized controlled trial of interventions for young people at ultra-high risk of psychosis: twelve-month outcome. J Clin Psychiatry 74:349–356.

McGrath J, Saha S, Chant D, Welham J (2008) Schizophrenia: a concise overview of incidence, prevalence, and mortality. Epidemiol Rev 30:67–76.

Menezes FP, Kist LW, Bogo MR, Bonan CD, Da Silva RS (2015) Evaluation of age-dependent response to NMDA receptor antagonism in zebrafish. Zebrafish 12:137–143.

Meyer U, Feldon J, Schedlowski M, Yee BK (2005) Towards an immuno-precipitated neurodevelopmental animal model of schizophrenia. Neurosci Biobehav Rev 29:913–947.

Meyer U, Nyffeler M, Schwendener S, Knuesel I, Yee BK, Feldon J (2008) Relative prenatal and postnatal maternal contributions to schizophrenia-related neurochemical dysfunction after in utero immune challenge. Neuropsychopharmacology 33:441–456.

Mezzomo NJ, Fontana BD, Müller TE, Duarte T, Quadros VA, Canzian J, Pompermaier A, Soares SM, Koakoski G, Loro VL, Rosemberg DB, Barcellos LJG (2019) Taurine modulates the stress response in zebrafish. Horm Behav 109:44–52.

Mezzomo NJ, Silveira A, Giuliani GS, Quadros VA, Rosemberg DB (2016) The role of taurine on anxiety-like behaviors in zebrafish: A comparative study using the novel tank and the light-dark tasks. Neurosci Lett 613:19–24.

Morales M, Spear LP (2014) The effects of an acute challenge with the NMDA receptor antagonists, MK-801, PEAQX, and ifenprodil, on social inhibition in adolescent and adult male rats. Psychopharmacology (Berl) 231:1797–1807.

Neuwirth LS et al. (2019) Assessing the Anxiolytic Properties of Taurine-Derived Compounds in Rats Following Developmental Lead Exposure: A Neurodevelopmental and Behavioral Pharmacological Pilot Study. Adv Exp Med Biol 1155:801–819.

O’Donnell CP, Allott KA, Murphy BP, Yuen HP, Proffitt T-M, Papas A, Moral J, Pham T, O’Regan MK, Phassouliotis C, Simpson R, McGorry PD (2016) Adjunctive Taurine in First-Episode Psychosis: A Phase 2, Double-Blind, Randomized, Placebo-Controlled Study. J Clin Psychiatry 77:e1610–e1617.

Oja SS, Saransaari P (1996) Taurine as osmoregulator and neuromodulator in the brain. Metab Brain Dis 11:153–164.

Oja SS, Saransaari P (2015) Open questions concerning taurine with emphasis on the brain. Adv Exp Med Biol 803:409–413.

Parksepp M, Leppik L, Koch K, Uppin K, Kangro R, Haring L, Vasar E, Zilmer M (2020) Metabolomics approach revealed robust changes in amino acid and biogenic amine signatures in patients with schizophrenia in the early course of the disease. Sci Rep 10:13983.

Powell SB, Geyer MA (2007) Overview of animal models of schizophrenia. Curr Protoc Neurosci Chapter 9:Unit 9.24.

Powell SB, Zhou X, Geyer MA (2009) Prepulse inhibition and genetic mouse models of schizophrenia. Behav Brain Res 204:282–294.

Rammes G, Mahal B, Putzke J, Parsons C, Spielmanns P, Pestel E, Spanagel R, Zieglgänsberger W, Schadrack J (2001) The anti-craving compound acamprosate acts as a weak NMDA-receptor antagonist, but modulates NMDA-receptor subunit expression similar to memantine and MK-801. Neuropharmacology 40:749–760.

Redmond HP, Stapleton PP, Neary P, Bouchier-Hayes D (1998) Immunonutrition: the role of taurine. Nutrition 14:599–604.

Salim S (2014) Oxidative stress and psychological disorders. Curr Neuropharmacol 12:140–147.

Saransaari P, Oja SS (2000) Taurine and neural cell damage. Amino Acids 19:509–526.

Schmack K, Bosc M, Ott T, Sturgill JF, Kepecs A (2021) Striatal dopamine mediates hallucination-like perception in mice. Science 372:eabf4740.

Seyhan AA (2019) Lost in translation: the valley of death across preclinical and clinical divide – identification of problems and overcoming obstacles. Translational Medicine Communications 4:18.

Shafer A, Dazzi F (2019) Meta-analysis of the positive and Negative Syndrome Scale (PANSS) factor structure. J Psychiatr Res 115:113–120.

Shirayama Y, Obata T, Matsuzawa D, Nonaka H, Kanazawa Y, Yoshitome E, Ikehira H, Hashimoto K, Iyo M (2010) Specific metabolites in the medial prefrontal cortex are associated with the neurocognitive deficits in schizophrenia: a preliminary study. Neuroimage 49:2783–2790.

Steullet P, Cabungcal J-H, Coyle J, Didriksen M, Gill K, Grace AA, Hensch TK, LaMantia A-S, Lindemann L, Maynard TM, Meyer U, Morishita H, O’Donnell P, Puhl M, Cuenod M, Do KQ (2017) Oxidative stress-driven parvalbumin interneuron impairment as a common mechanism in models of schizophrenia. Mol Psychiatry 22:936–943.

Stone JM, Morrison PD, Pilowsky LS (2007) Glutamate and dopamine dysregulation in schizophrenia--a synthesis and selective review. J Psychopharmacol 21:440–452.

Stubbs B, Williams J, Gaughran F, Craig T (2016) How sedentary are people with psychosis? A systematic review and meta-analysis. Schizophr Res 171:103–109.

Tran S, Muraleetharan A, Fulcher N, Chatterjee D, Gerlai R (2016) MK-801 increases locomotor activity in a context-dependent manner in zebrafish. Behav Brain Res 296:26–29.

Volk DW, Lewis DA (2014) Early developmental disturbances of cortical inhibitory neurons: contribution to cognitive deficits in schizophrenia. Schizophr Bull 40:952–957.

Winter C, Djodari-Irani A, Sohr R, Morgenstern R, Feldon J, Juckel G, Meyer U (2009) Prenatal immune activation leads to multiple changes in basal neurotransmitter levels in the adult brain: implications for brain disorders of neurodevelopmental origin such as schizophrenia. Int J Neuropsychopharmacol 12:513–524.

Woods S, Saksa J, Compton M, Daley M, Rajarethinam R, Graham K, Breitborde N, Cahill J, Srihari V, Perkins D, Bearden C, Cannon T, Walker E, McGlashan T (2017) 112. Effects of Ziprasidone Versus Placebo in Patients at Clinical High Risk for Psychosis. Schizophr Bull 43:S58.

Yang J, Guo H, Sun D, Duan J, Rao X, Xu F, Manyande A, Tang Y, Wang J, Wang F (2019) Elevated glutamate, glutamine and GABA levels and reduced taurine level in a schizophrenia model using an in vitro proton nuclear magnetic resonance method. Am J Transl Res 11:5919–5931.

Zanos P et al. (2016) NMDAR inhibition-independent antidepressant actions of ketamine metabolites. Nature 533:481–486.

Zoicas I, Kornhuber J (2019) The Role of the N-Methyl-D-Aspartate Receptors in Social Behavior in Rodents. Int J Mol Sci 20:E5599.

